# Persistent declines in forest-dependent birds following active restoration of logged tropical forest in Borneo

**DOI:** 10.64898/2026.02.15.705981

**Authors:** Gianluca Cerullo, Andrew Balmford, Suzan Benedick, Catherine Finlayson, Toby Jackson, Tommaso Jucker, Daniel Kong, Simon C. Mills, Simon Mitchell, Oscar Morton, David P. Edwards

**Affiliations:** Department of Zoology, University of Cambridge, Cambridge, UK; Conservation Research Institute, University of Cambridge, Cambridge, UK; Forest Ecosystems & Society, Oregon State University, Corvallis, OR, United States; Faculty of Sustainable Agriculture, Universiti Malaysia Sabah, Locked Bag No. 3, 90509, Sandakan, Sabah, Malaysia; Center for Natural Climate Solutions, Conservation International, Arlington, VA, USA; School of Biological Sciences, University of Bristol, Bristol, UK; Malaysian Nature Society Kuching Branch, P.O. Box A144, Kenyalang Park, 93824, Kuching, Sarawak, Malaysia; Rainforest Builder, London, UK; Durrell Institute for Conservation & Ecology, School of Anthropology and Conservation, University of Kent, Canterbury, UK; Ecology and Evolutionary Biology, University of Sheffield, UK; Department of Plant Sciences and Centre for Global Wood Security, University of Cambridge, Cambridge, UK

## Abstract

1. Tropical forest restoration is critical for mitigating biodiversity loss and climate change, including in forests impacted by selective logging. Active restoration through liana cutting and enrichment tree planting can substantially accelerate carbon recovery, potentially reducing economic pressures to convert logged forests. But its long-term biodiversity impacts remain largely unknown.
2. Using over two decades of bird survey data from Borneo’s largest logged-forest restoration project, we quantified occupancy patterns for 176 species across primary, naturally regenerating, and actively restored logged forests spanning a 30+ year post-logging chronosequence.
3. Forest-dependent, threatened and near-threatened species generally declined through time in actively restored areas, whereas many species in naturally regenerating forests progressively recovered toward primary forest levels. Between 17–40% of 66 threatened or near-threatened species had consistently lower occupancies in actively restored than in naturally regenerating forest. Across species of global conservation concern, median occupancies in restored areas remained ∼22% below primary forest even 50 years after harvests, compared with only ∼6% lower under natural regeneration.
4. Arboreal insectivores, frugivores, and predatory species appeared most negatively affected by active restoration, with 27-49% of arboreal gleaning insectivores (of 62), 13-30% of arboreal frugivores (of 40), and one-third of predatory species (of 15) showing higher occupancy in naturally regenerating forests. Sallying insectivores also showed a possible but uncertain response, whereas ground-associated frugivores and insectivores were largely unaffected by restoration treatment.
5. Concerningly, even 50 years post-logging, up to 52% of 50 high forest-dependency species retained distinct occupancies in actively restored compared with primary forest, suggesting persistent negative impacts of vine-cutting and/or tree planting activities on avian populations.
6. **Synthesis and applications**. Our findings indicate that despite substantial carbon benefits, active restoration within selectively logged forests may impede the recovery of forest-dependent biodiversity. This challenges the common assumption embedded within nature-based climate solutions that carbon and biodiversity outcomes will necessarily align. Nonetheless, despite the persistent declines in bird communities, actively restored forests continued to provide key habitat for many species. Active interventions may thus still contribute to broader biodiversity conservation objectives if they protect logged areas from conversion, potentially via carbon payments.

## Introduction

Tropical forest restoration is accelerating globally, driven by major commitments to mitigate climate change and biodiversity loss^1,2^. The Bonn Challenge aims to restore 350 million hectares (Mha) by 2030, while the Kunming-Montreal Global Biodiversity Framework commits to conserving and recovering 30% of the world’s lands^3^. To date, most political and scientific attention has focused on the potential ecological and climate gains from enabling secondary forest regeneration on former or current agricultural lands^1,4,5^, showing that forest regrowth in priority regions can cost-effectively reduce extinctions and sequester ∼23 Gt C over the next three decades ^5^. However, large-scale restoration of agricultural areas carries the risk of displacing food production into remaining intact forests, potentially undermining both biodiversity and climate goals^2,6^. In contrast, relatively little research has explored the long-term ecological impacts of restoring degraded tropical forests that have had persistent forest cover ^7^. This is despite growing evidence that a substantial portion of global forest restoration potential lies within existing degraded forestlands^8,9^, where interventions also carry notably lower risks of agricultural displacement.

More than 400 million hectares of tropical forest are designated for timber production and have been, or soon will be, selectively logged, involving the removal of large merchantable trees while maintaining partial canopy cover; a large additional area is also affected by illegal harvests. Selective logging structurally alters tropical forests, causes the reduction of forest-specialist biodiversity, and drives short-term losses of carbon storage and other ecosystem services. Yet logged forests represent a critical land base that could contribute to ambitious national restoration commitments^7,12,13^. Several restoration approaches exist within logged landscapes. Passive natural regeneration involves removing additional pressures following logging to enable forests to recover without assistance, and is the most common approach pantropically^10^. However, forest managers are increasingly using active interventions, especially in heavily degraded areas where natural regeneration potential is impaired or where accelerating aboveground biomass or merchantable timber stock recovery is a primary goal^7,14^. Key active interventions include the cutting of lianas and other climbers which proliferate in the lighter post-harvest understorey, as well as the enrichment planting of overexploited tree species^7,15^. Both actions have relatively well-evidenced climate and timber recovery benefits compared to passive regeneration^13,15–17^. For example, one pantropical meta-analysis found that liana cutting was associated with an 88% increase in carbon accumulation relative to untreated forests^18^, while enrichment planting more than doubled the recovery rate of merchantable timber in lowland dipterocarp forests in Indonesia^17^. Where intensive logging has left vast areas of degraded forest with declining timber yields and high agricultural conversion pressures (e.g., especially in SE Asia), active restoration of logged forests is thus increasingly promoted as a nature-based climate solution^12,19^.

As restoration agendas gather momentum, there is growing expectation that forest restoration should deliver biodiversity co-benefits alongside any climate gains^20,21^. Yet the long-term, species-level impacts of restoration in selectively logged tropical forests remain largely unknown. Unlike reforestation on former agricultural land, where biodiversity recovery typically follows predictable increases in recolonization by forest-using species and eventual community convergence towards reference compositions^22^, selective logging leaves behind a complex mosaic of mostly closed-canopy patches harvested at different intensities and times and interspersed with unlogged patches^23^, resulting in highly variable regeneration potential. Within this context, active restoration via liana cutting and enrichment planting may accelerate the recovery of old-growth structures and resources, thereby benefiting forest specialist species (e.g. by increasing coverage of large trees). However, such interventions could also reduce resource availability and heterogeneity of recovering forests, especially because lianas are ecologically important as nesting and foraging substrates, and because planting often emphasizes a narrow range of merchantable timber species^7^. Detecting whether active restoration confers biodiversity benefits beyond passive regeneration in logged forests is therefore important but also challenging because it requires field data tracking species-level responses across a wide range of post-harvest recovery times and restoration types, plus unlogged reference sites. Yet all studies exploring the biodiversity impacts of logged forest restoration to date have focused on short-term impacts^7,19,27–29^, overlooking the temporal and species-specific dynamics critical for designing effective, lasting restoration strategies.

Here, we overcome these challenges using more than two decades of systematically collected avian data from Malaysian Borneo, the site of the tropics’ largest and longest-running logged forest restoration project. Our sampling effort includes 10,368 detections of 176 bird species across 200 survey points spanning old-growth, naturally regenerating, and actively restored once-logged forests (subject to liana cutting and enrichment planting), surveyed along a >30-year post-logging chronosequence. Research in our study landscape has previously shown that compared to natural regeneration, restoration increased annual aboveground biomass accumulation by 50% in the three decades following logging^12^, and that short-term outcomes for birds following restoration included the retention of insectivorous species but losses in frugivores and phylogenetic diversity^27,28^. We ask two key unresolved questions: (i) how do long-term restoration outcomes vary with species’ ecology and conservation status; and (ii) does active restoration enhance biodiversity recovery in logged forests over time compared with natural regeneration?

## Methods

### Study site

Avian surveys were carried out in the 1Mha Yayasan Sabah concession, among the largest remaining contiguous blocks of lowland dipterocarp rainforest in Southeast Asia, encompassing a mixture of primary and selectively logged forests (Figure 1)^30^. Primary forest sites were located within the Danum Valley Conservation Area (DVCA; 43,800 ha, 80 points), with one additional site (12 points) located in the nearby Tawau Hills Park (28,000 ha). Logged forest sites were within the Ulu Segama Forest Reserve (USFR; 238,000, 108 points), adjacent to Danum Valley. The USFR was selectively logged once between 1972 and 1993, extracting an average of 113 m^3^ ha^−1^ of timber, with some remnant patches of unlogged forest remaining in steep or riparian zones^30,31^. Naturally regenerating portions of this landscape contain our ‘once-logged’ naturally regenerating sites. A section of once-logged forest was actively managed under the INFRAPRO Forest Rehabilitation Project^12,27^. This large-scale restoration project began in the early 1990s with the goal of accelerating the recovery of forest structure and carbon. Restoration treatments were applied to once-logged areas averaging nine years post-harvest, using remaining logging infrastructure for planting and cutting access. Treatments included liana removal, line planting of over 50 native dipterocarp and fruit tree species, and maintenance of open planting strips through repeated liana cutting events over three years (see Philipson et al. 2020 for details^12^). Between 1993 and 2004, INFRAPRO restored patches ranging from 83 to 1,745 ha – our ‘restored’ sites all fall within these treated areas (Figure 1).

**Figure 1.**
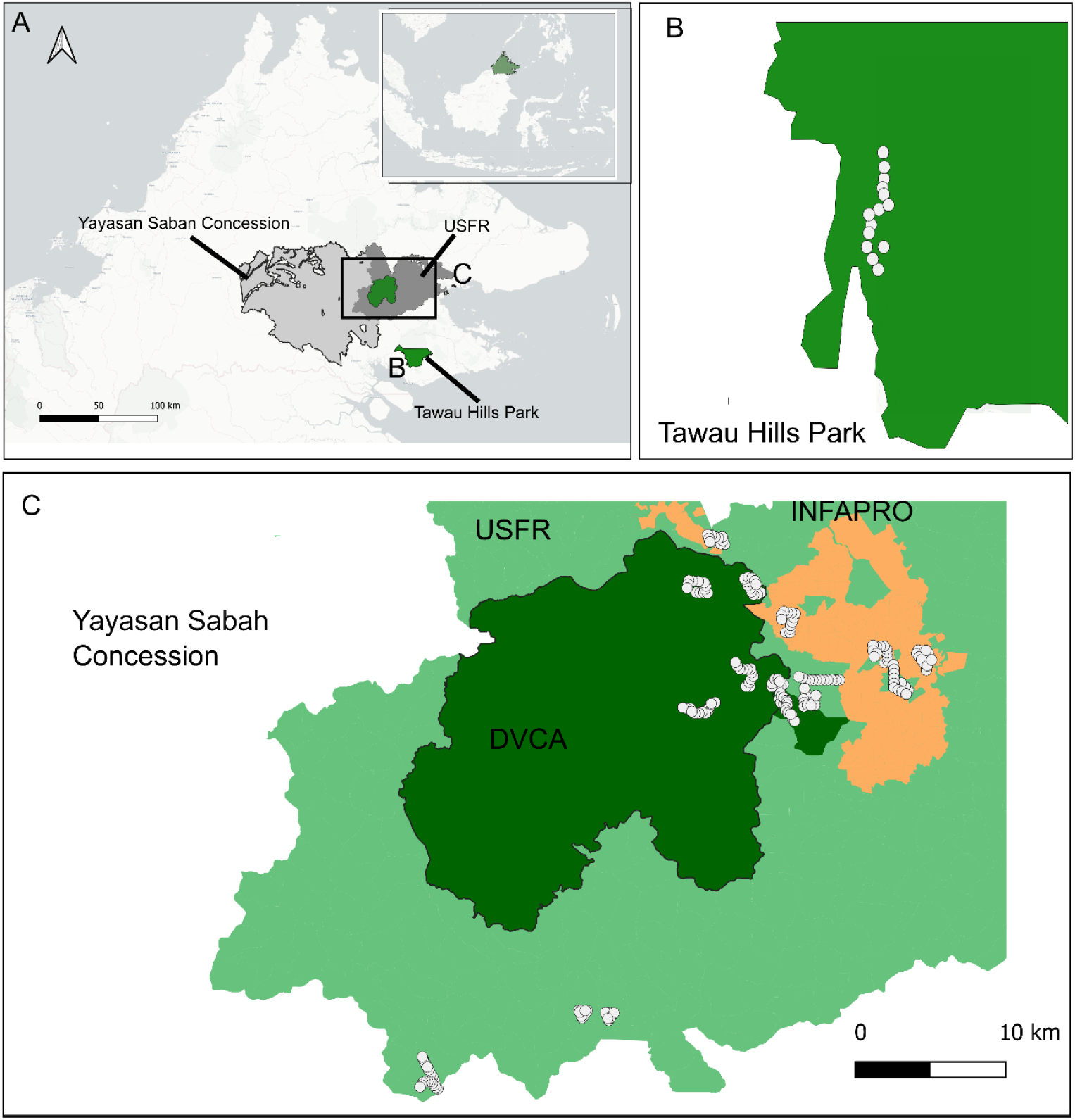
Location of sample points in Sabah Malaysia. Avian point counts were located in the Yayasan Sabah logging concessions, predominantly with the Ulu-Segama Forest Reserve (A), with a small number of primary forest points located in Tawau Hills Park (B). Points within the Danum Valley Conservation Area (Dark Green) were primary forest, and our once-logged points were in the USFR and wider Yayasan Sabah area. A section of this once-logged forest was actively restored as part of the INFAPRO restoration area (yellow), with remaining once-logged forest left naturally regenerating (light green).

### Bird Surveys

Avian point counts were conducted following Edwards et. 2011 and Mitchell et al. 2018, covering 200 point-locations and 10,368 detections of 172 species, excluding flyovers. Briefly, sites were established at least 500 m apart and 200 metres from roads or edges, with sites consisting of transects of 9–12 point-locations separated by >200 m to maintain independence. We returned to survey 90% (n=180) of points over 3 consecutive days, while 10% of points were surveyed across 4 days. Points were stratified across primary forest (8 sites, 92 points), naturally regenerating once-logged forest (5 sites, 60 points; post-harvest range = 19-62 yrs) and restored forests (4 sites, 48 points; post-harvest range = 21-41 yrs). At each point, all birds seen or heard within 100 m of the point locations were identified, with surveys conducted on mornings with no rain between 05:50 and 11:00 between May and Aug in 2008, 2009, 2011, 2017 and 2022 by three expert ornithologists (DE, SM, DK).

### Occupancy-detection modelling

#### Occupancy

To examine how alternative restoration approaches affected bird species occupancy through time, while accounting for imperfect detection, we fitted a hierarchical Bayesian multispecies detection-occupancy model^34,35^. This allowed us to estimate species-specific responses to restoration treatments, plus primary forest reference conditions, while accounting for variables influencing detection. *A priori*, we expected that species occupancy would vary according to the management treatments (‘primary’, ‘once-logged’, ‘restored’), as well as the time since logging, and that these changes would interact with species’ habitat preferences (particularly their degree of forest dependency). To capture these sources of variation, we classified each bird species detected in field surveys as having either “high”, “medium” or “low”, forest dependency (50, 107, 18 species, respectively), following Birdlife International’s classifications **(REF)**, but merging ‘low’ and ‘not normally found in forest’ categories, due to low sample sizes^36^. We then pooled variance within these categories, assuming that species in the same category responded on average more similarly to forest management. Such pooling improves occupancy estimates for rarely detected species, which we expected to be most sensitive to forest structural change^37^.

Specifically, we structured the occupancy component of our models as:

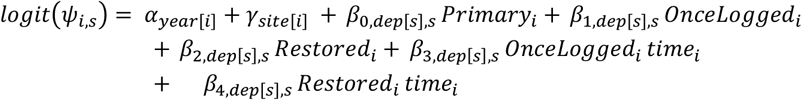

Here, Ψ_*i,s*_ is the probability that species, *s*, is present at point *i*. Any community-wide annual variation in occupancy is accommodated by a random intercept on year, *α* _*year* [*i*]_, where *year*[*i*] refers to the sampling year of point *i*; and spatial variation by a random intercept for site, *γ*_site[*i*]_. Habitat-effects are captured using a series of mutually exclusive habitat dummy variables (i.e. a point can only be one habitat type). The subscripting on each parameter indicates species-specific random slopes for forest treatment and time-since-logging effects (denoted by dep[s], which identifies the forest-dependency class of species *s* and within which the species-specific effects are partially pooled). Thus, this structure allowed each species to have its own effect (slope) for each forest treatment and its interaction with time since logging, while borrowing strength from other species within the same hierarchical-dependency class. *time[i]* represented the time since logging at point *i*, included in interaction terms to capture temporal dynamics in occupancy recovery within naturally regenerating and restored logged forests.

#### Detection

The detection component was modelled as a function of management treatment, forest structure, time of day, and point-count observer identity, while also capturing species-level variation in detectability. Fixed effects included the management treatment (primary, once-logged, restored), forest structure within a 50 m radius buffer, calculated for each point from an above-ground density LiDAR product^38^ (ABD50; see below), time of day, and observer. To account for variation in detection across observers and species, we included a random intercept for each observer–species combination. We also allowed detection to vary across species as a function of time of day by including a species-specific random effect.

Specifically, detection was modelled as:

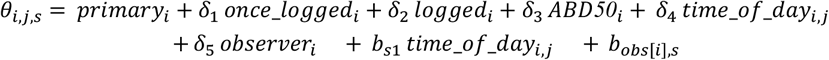

Here, *θ*_*i,j,s*_ represented the probability of detecting species *s* during visit *j* at point *i*. The term *observer*_*i*_ refers to the observer (DE, SM, DK) conducting the survey at point *i*, and *b*_*s*_ *time_of_day*_*i,j*_ captures the specific time of the *j*-th visit at point *i*. The random effect *b*_*s*1_ modelled species-specific variation in how detectability changes across the day, while *b*_*obs*_[_*i*_]_,*s*_ accounted for variation in detection across observer–species pairings. To capture differences in vegetation structure that might have affected visibility at point count locations, we included ABD50, calculated as the mean above-ground density within a 50 m-radius buffer of each survey point. Density was derived from a 30 m resolution LiDAR product for the year 2016, which falls approximately midway between the earliest and latest years of bird sampling (2008–2022)^38^. We fitted the occupancy detection model in Stan (v2.26.1) via cmdstanr (v0.5.3), using the flocker package (v0.5.3)^35^. All models were run with four chains of 2000 warm-up and 4000 post-warm-up iterations. We applied weakly informative priors on all parameters and monitored convergence through standard diagnostics with model adequacy evaluated via posterior predictive checks.

### Threat status and guild data

We examined species-level responses as a function of (i) threat status, derived from the IUCN Red List (Critically Endangered [CR], Endangered [EN], Vulnerable [VU], Near Threatened [NT] and Least Concern [LC]; (ii) species forest dependency (described above); and (iii) foraging-dietary guilds. Species were assigned into separate foraging guilds (i.e. arboreal, sallying and terrestrial foragers, the latter including undergrowth foragers) and feeding guilds (i.e. frugivores, insectivores, predators, nectarivores, or piscivores), following Edwards et al 2012. These guilds capture differences in resource use and forest strata and may therefore influence responses to restoration; however, we pooled variance by forest dependency to impose a weakly informative hierarchical structure, reflecting prior expectations that species respond coherently to forest structure and disturbance intensity. To assess uncertainty in the differences in species occupancy between restored and once-logged forests, we calculated the *Probability of Direction* (or pd), which is defined as the proportion of posterior samples that share the same sign as the median estimate (either positive or negative)^39^. Thus, a pd of 0.5 indicates no consistent directional effect (i.e., equal support for higher occupancy in either habitat), whereas a pd of 1 indicates complete agreement among posterior draws in the direction of the effect (e.g., consistently higher occupancy in naturally regenerating than restored forests, or vice-versa). We applied probability of direction (pd) thresholds of 0.75 and 0.9 to represent less and more conservative evidence cutoffs, respectively, reflecting the need in conservation practice to balance sensitivity to potential forest management interventions with caution against over-interpreting uncertain trends. We used these same thresholds to evaluate temporal consistency in species’ responses by assessing whether changes in occupancy between different time points (e.g., early vs. late recovery stages) showed a consistent direction across posterior draws. This allowed us to identify which species had robust directional trends through time.

## Results

### Impacts by species global conservation status

We detected 66 globally threatened or near-threatened species (CR, EN, VU, NT) across the landscape. Of these birds of conservation concern, over 40% (28 species) showed consistent differences in occupancy between restored and once-logged forests (at pd > 0.75), decreasing to ∼17% of species (*n =* 11 species) when using a more stringent cutoff of *pd* > 0.9 (Figure 2). Of these potentially affected species, 92-100% (26 of 28 species where pd > 0.75 and 11 of 11 where pd > 0.9) had higher occupancy in once-logged than actively restored forest, including species like Rufous-crowned Babbler (NT; *pd* = 0.91), Yellow-crowned Barbet (NT; *pd* = 0.91), and Scarlet-rumped Trogon (NT; *pd* = 0.94). Only two species of conservation concern, Greater Green Leafbird (EN; *pd* = 0.80) and Chestnut-necklaced Partridge (VU; *pd* = 0.80), showed a consistently higher predicted occupancy in actively restored forests than in naturally regenerating forest (Figure S1 shows species-level estimates with associated uncertainties). For all birds of conservation concern, median occupancies in actively restored forest were ∼22% below primary forest even after 50 years of recovery, whereas in naturally regenerating forests occupancies were only modestly lower (6%) than in primary forest. Among the 106 least-concern species (Figure S2), 16–64% (16–33 species) showed higher occupancy in naturally regenerating forest, whereas only 15% (16 species) did so in actively restored stands at pd > 0.75, and none at pd > 0.9. Despite this, least-concern species showed relatively small absolute differences in median occupancy compared to primary, with median occupancies only ∼2% below primary-forest levels in actively restored, compared with ∼10% lower in naturally regenerating forest.

**Figure 2.**
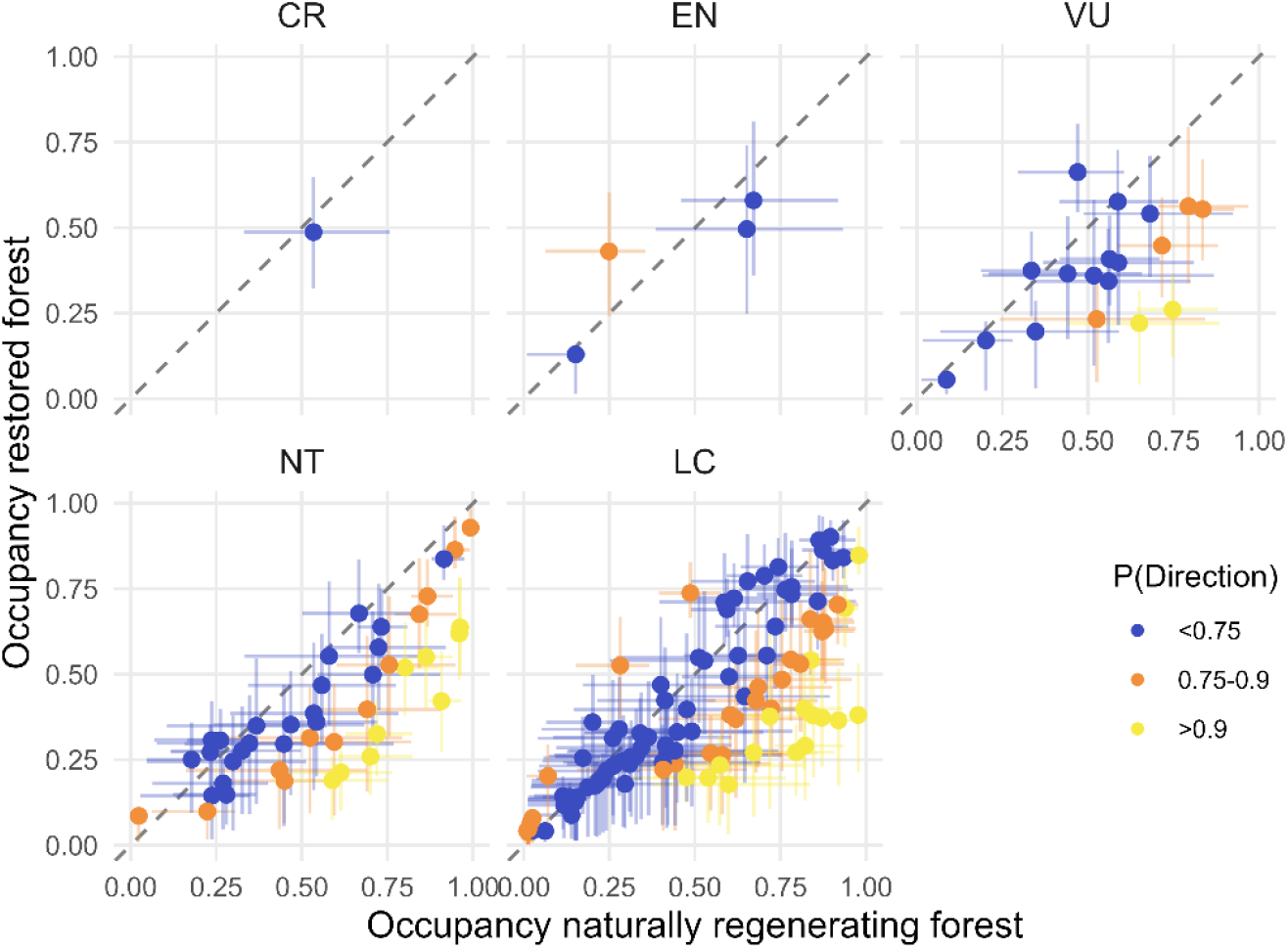
Relationship between estimated avian occupancy in naturally regenerating once-logged and actively restored forests across a lowland dipterocarp-dominated rainforest landscape in Sabah, Borneo. Each point represents the mean occupancy for a species, faceted by IUCN Red List threat status: Critically Endangered (CR), Endangered (EN), Vulnerable (VU), Near Threatened (NT), and Least Concern (LC). Horizontal and vertical bars represent 50% (solid) credible intervals for occupancy estimates calculated across 500 posterior draws per year between 19–50 years post-logging. The 1:1 line (dashed) indicates equal occupancy between forest types. Points above the line reflect species with higher mean occupancy in restored than once-logged forests, and vice versa. Points are coloured by the *probability of direction* (PD), which reflects the consistency in the direction of the restoration effect across posterior draws. Colours indicate greater certainty that occupancy differs between restoration treatments. Outcomes by forest dependency class are in Figure S4.

### Impacts across feeding-foraging guilds and forest dependency

While species from most feeding–foraging guilds showed some response to restoration treatment, a greater proportion of arboreal and predatory species were affected, and/or their responses were more consistently detected (Figure 3). Nearly half of arboreal gleaning insectivores had higher occupancies in naturally regenerating than in actively restored forest (30/62 species at pd > 0.75, declining to 17/62 at pd > 0.9; Figure 3), and median occupancies in restored forest were ∼12% lower than in primary baselines (versus 17% higher under natural regeneration). Arboreal frugivores and mixed frugivore– insectivores showed similar patterns: 30% (12/40 species) favoured naturally regenerating forest at pd > 0.75, falling to ∼13% (5/40) at pd > 0.9. This group included several species with apparent preferences for naturally regenerating forest, including Green Broadbill (pd = 0.86) and Blue-rumped Parrot (pd = 0.80). Indeed, arboreal frugivores’ median occupancies were ∼10% higher than primary forest in naturally regenerating stands but moderately (∼4%) lower in actively restored forest. The minority of frugivores which were more common in restored forest including Olive-winged Bulbul (pd = 0.76), Yellow-vented Bulbul (pd = 0.77), and Yellow-breasted Flowerpecker (pd = 0.83). Predators also showed clear impacts from restoration treatments, with 33% (5/15 species) having higher occupancy in naturally regenerating than restored forest at pd > 0.75, falling to 20% (3/15) at pd > 0.9. These included large-bodied species such as Bushy-crested Hornbill (pd = 0.85) and Rhinoceros Hornbill (pd = 0.86) and there was a general tendency for larger-bodied birds to prefer naturally regenerating forest (Figure S3). Median predator occupancies were ∼20% lower in actively restored compared to primary forest (*cf*. only 4% lower with natural regeneration).

**Figure 3.**
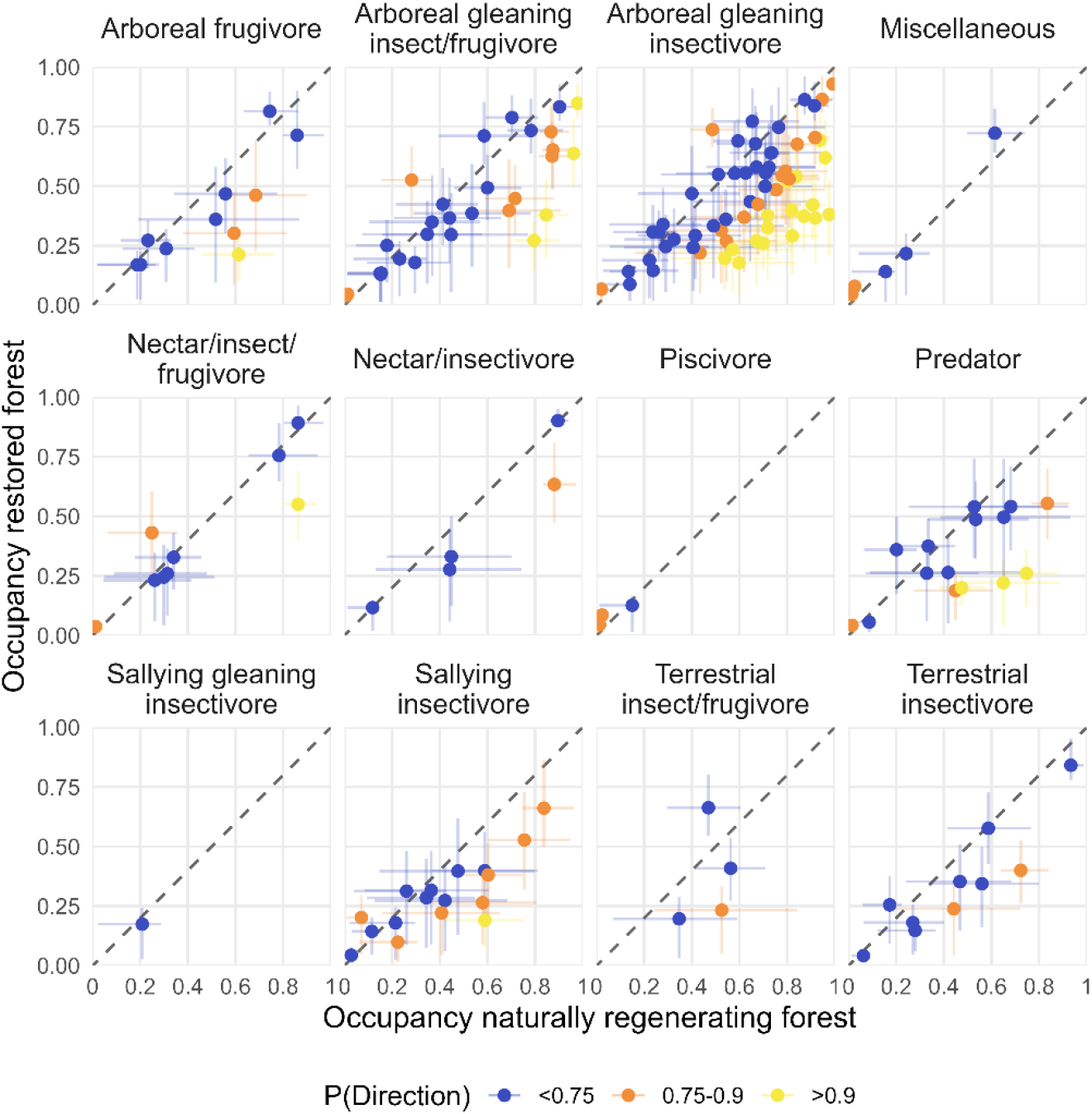
Relationship between species’ estimated occupancy in restored and once-logged forests, grouped by foraging-feeding guild. As in Figure 2, each point represents the mean occupancy for a species in restored (y-axis) and naturally regenerating once-logged (x-axis) forests, with credible intervals showing 50% credible intervals calculated across 500 posterior draws per year between 19– 50 years post-logging. Colours indicate greater certainty that occupancy differs between restoration treatments. Outcomes by forest dependency class are in Figure S4.

Sallying insectivores also appeared sensitive to restoration actions, but this was more uncertain. At a lower evidence threshold (pd > 0.75), 41% of species (7/17) had higher occupancy in naturally regenerating forest, but this dropped to just one species—the Scarlet-rumped Trogon (pd = 0.9)—under a stricter cutoff (pd > 0.9). Across the guild, median occupancies were ∼13% lower than primary forest in restored stands, compared to ∼5% lower with natural regeneration. By contrast, more ground-associated birds, including terrestrial insectivores and mixed terrestrial insectivore–frugivores, showed limited and uncertain responses. Only 21% of species belonging to these guilds (3/14) crossed pd > 0.75, and none met pd > 0.9. The few species showing consistent differences at lower thresholds included Black-hooded Pitta (pd = 0.86) and Hooded Pitta (pd = 0.83), both favouring natural regeneration. Yet even though effects were uncertain for terrestrial species, median occupancies were still ∼28% lower than primary, compared to only ∼4% lower in natural regeneration, reinforcing the overall pattern of reduced recovery in restored stands. These trends persisted when examining species by forest-dependency class (Figure S4-6). Among high-dependency species, 28–48% (14–24 of 50 species, depending on the pd threshold) showed higher occupancy in naturally regenerating forest, while medium-dependency species exhibited similar but weaker patterns, with 12–31% (13–33 of 107 species) favouring naturally regenerating over restored forests.

### Temporal recovery within restoration treatments

Species’ post-logging recovery trajectories differed between restoration treatments and were influenced by forest dependency, threat status, and foraging guild (Figure 4, Figure S7–S8). High forest-dependency species showed the clearest long-term gains under natural regeneration: after ∼50 years, median occupancy in naturally regenerating forest averaged 10% higher than in primary forest, and 96% (48/50) of species increased between 20- and 50-years post-logging. These positive trends were consistent for 59% (28 species at pd > 0.75) and 23% (11 species at pd > 0.9), with nearly half of these showing no detectable difference from primary-forest occupancy, indicating recovery to pre-logging levels. Recovering species included Gold-whiskered and Yellow-crowned Barbets and Red-naped and Scarlet-rumped Trogons. In contrast, only 4% (2/50) of high-dependency species increased through time in restored stands. Median occupancy in restored forest harvest 50 year previously remained 31% below primary levels, and 52% (26 species, pd > 0.75) retained occupancy deficits after five decades. This fell to 12% (6 species; pd > 0.9) under a more conservative threshold, highlighting both the uncertainty around recovery trajectories in restored forest and the potential for actively restored stands to continue supporting avifaunal communities.

**Figure 4.**
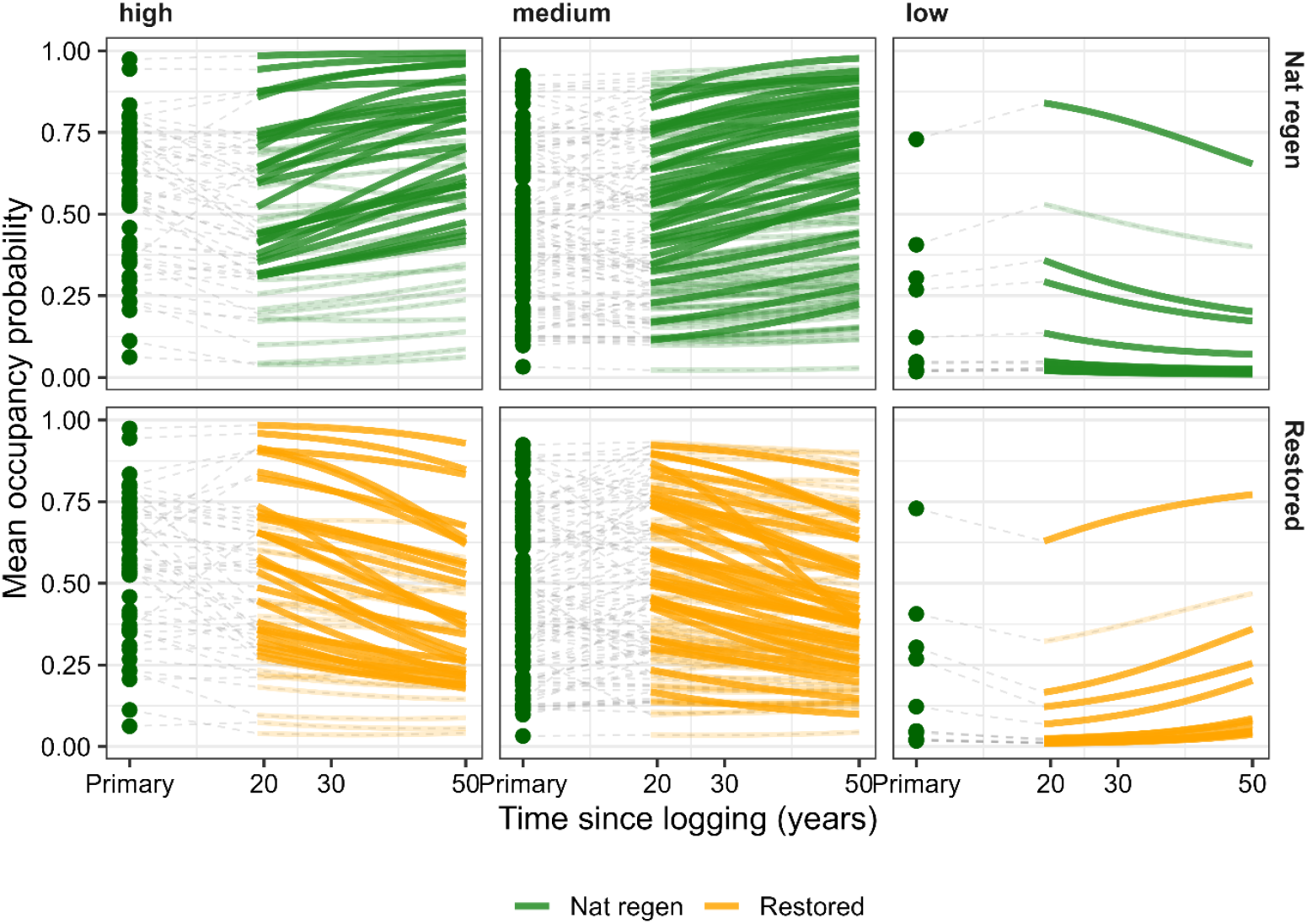
Temporal trends in species’ mean occupancy probability in once-logged (top row) and restored (bottom row) forests, and in primary forest, across a lowland dipterocarp rainforest landscape in Sabah, Borneo. Green dots represent a given species’ mean occupancy in primary forest, while each solid line represents a species’ mean occupancy in a given management treatment between 19–50 years post-logging, calculated across 500 posterior draws per year. Columns represent species grouped by their overall forest dependency: high, medium, and low, as defined by Birdlife International. Grey dashed lines represent the period (0-19 years post-logging) where we lack sampling coverage and therefore do not make predictions of occupancy. Temporal outcomes by foraging guild and threat status are in Figure S7-S8. Bold lines have a consistent directional difference in occupancy between 19 and 50 years post-logging within a particular restoration treatment (pd > 0.75).

Overall, these temporal patterns were similar for medium forest-dependent species (Figure 4 B,E; FigureS5). Of 107 species, 86% (92 species) increased in median predicted occupancy in naturally regenerating forests, with trends consistent for 40% (37 species at pd >0.75) and 7% (8 species at pd >0.9), and with median occupancies on average ∼20% higher in naturally regenerating versus primary forest. In actively restored forest, ∼8% (8 species) of medium forest-dependent species retained distinct occupancies from primary forest 50-years post-logging, although absolute differences were small and more similar to primary forest (∼3% below primary). Patterns for the small number of low forest-dependency species detected were reversed, with median occupancy declining under natural regeneration but increasing modestly in restored stands (Figure 4 C,F).

Threatened species also recovered primarily in naturally regenerating forest (Figure S7). Of 66 globally threatened or near-threatened species, 89% (59 species) increased between 19 and 50 years after logging, with consistent increases for 9–41% (6–37 species across pd thresholds). By contrast, only 9% (6 species) increased in actively restored forest, with only a single species, the Helmet Hornbill, showing consistent trend above pd >0.9.

Recovery trajectories also varied across foraging guilds (Figure S8). Arboreal guilds showed the clearest recovery with natural regeneration, with 17–41% (16-39 species for respective thresholds) of arboreal insectivores and frugivores increasing through time. Terrestrial insectivores (9–36%; 1-4 species) and predators (6–37%; 1-6 species) showed similar but slightly weaker positive trends, while sallying insectivores showed uncertain recovery trajectories: 41% (7 species) increased at pd > 0.75, but none at pd > 0.9. In contrast, recovery in actively restored forest was limited across all guilds, with only 3% (2 species) of arboreal gleaning insectivores, 6% of sallying insectivores (1 species), 6% (1 species) of predators, and 25% (1 species) of terrestrial insectivore–frugivores showed increasing at pd > 0.75; again, only the Helmeted Hornbill met pd > 0.9.

## Discussion

Our multi-decade, species-level analysis provides the first broad-scale, long-term evaluation of how the biodiversity gains of active restoration in selectively logged tropical forests compare with those of natural regeneration^7^. Although research in our study landscape previously revealed that enrichment planting and vine-cutting increased rates of annual carbon recovery by ∼50%, offering potentially cost-effective carbon drawdowns^12^, we found little evidence that these benefits have been complemented by increases in avian occupancy. Instead, forest-dependent and threatened species have generally declined through time in actively restored areas, whereas many species in naturally regenerating logged forests have gradually recovered towards primary forest levels. Despite uncertainty around species-level responses in our estimates – likely driven by high spatial and temporal variability in forest structure and resource availability within selectively logged forests^10,23,40,41^ – our findings challenge the common assumption embedded with nature-based climate solutions that carbon and biodiversity benefits will necessarily align^22,42,43^.

The apparently persistent declines we observed in threatened and high forest-dependency species likely result from the long-term effects of restoration interventions on forest structure and composition. Liana removal is designed to release regrowing trees from competition and accelerate canopy recovery^16,18^, and repeat-LiDAR surveys in our wider landscape indicate that vine-cutting enhances growth, reduces mortality of large trees, and accelerates post-logging canopy gap closure (Jackson *et al*., *in review*). However, vine cutting also removes important nesting and foraging substrates, fruit resources, and movement pathways for wildlife, with these features being more likely to persist or recover in naturally regenerating forest^7,27,44,45^. This may explain the more pronounced declines we observed in arboreal insectivores and frugivores, as well as large-bodied predatory species in actively restored forests: species in these guilds rely on more heterogeneous canopy structures, fruits, or invertebrate or vertebrate prey resources hosted within vine tangles, that may be lost after vine cutting or in the denser or darker canopies recovering of actively restored forests^25,46^. By contrast, sallying and particularly more-terrestrial species showed less clearcut differences in their responses to active and passive restoration, potentially because these species’ prey base (ground leaf-litter prey or aerial insects) is less associated with forest structural attributes. Enrichment planting in our study also incorporated a high diversity of timber species (up to 50 species), so it is possible that understorey microclimates and ground-level conditions were relatively comparable in naturally regenerating and actively restored forests^15,29^.

The sustained apparent occupancy losses among large-bodied frugivores and threatened or near-threatened species is particularly concerning because it implies erosion of key ecosystem functions and potentially elevated extirpation risks following active restoration^46,47^. This aligns with previous findings that actively restored forests support avian communities with reduced functional and phylogenetic diversity^48^. Our results extend these insights by revealing the temporal dynamics of species’ responses, showing that many forest-dependent species continue to decline many decades after logging and restoration interventions have ceased, whereas naturally regenerating forests tend to recover gradually towards primary forest occupancy levels. Although the relatively high uncertainty around these contrasting trajectories underscores the value of additional future sampling campaigns within recovering logged forests, these patterns could be explained by short-term demographic bottlenecks in the aftermath of multiple rounds of liana cutting leading to long-term declines and extinction debts, or else be driven by active interventions leaving lasting structural and prey-based legacies that constrain bird recovery^49^. Our findings thus highlight the need for greater investment in long-term biodiversity monitoring as well as future studies that track abundance and demography directly^28^, ideally including immediate post-cutting surveys and before–after control–impact designs to overcome the limits of space-for-time approaches^50,51^. Future studies should also explore restoration impacts on other taxa and in more heavily degraded forests than those in our study landscape, since active interventions could still provide benefits not detected here, particularly where repeated logging has more seriously impaired natural regeneration potential via intensified liana infestation^52,53^.

Although it was not possible to disentangle the effects of liana cutting from enrichment planting in our study landscape, the financial costs of enrichment planting alone are substantial (≈$1500-$2500 USD ha^−1^ in the project area^12^). These high costs, combined with the stronger recovery of forest-dependent and threatened species in naturally regenerating forests, suggest that when biodiversity recovery is the primary goal of restoration, investments may be better directed towards supporting natural regeneration in areas of high recovery potential, such as through targeted purchases of logged-forest land or community stewardship programmes^31,54^. Modifying the active restoration treatment (e.g. via more-targeted liana cutting or a reduced emphasis on strip-planting) could reduce active restoration costs and presumably also the apparent trade-off between carbon and biodiversity outcomes. However, given rampant avian trapping and hunting pressures across Southeast Asia, funding community-led patrols and anti-hunting initiatives in logged-over areas may ultimately provide more cost-effective and reliable returns for biodiversity than expensive silvicultural interventions focused primarily on timber or carbon recovery^55–57^ Nevertheless, the reality in much of Southeast Asia is that many logged-over forests lie within inactive concessions, have been stripped of much of their commercial timber, and now face an increased risk of forest conversion to industrial plantations, with disastrous consequences for biodiversity^11,32,58–60^. Where active restoration can prevent such conversion by extending forest protection across restored logged forests, potentially supported by carbon finance, it may still deliver substantial biodiversity benefits relative to conversion. Indeed, we show that actively restored forests do appear to continue to provide habitat for many species, including threatened. However, where the realistic counterfactual to active restoration is natural regeneration, our findings indicate that passive recovery may offer superior outcomes for biodiversity, even if not for carbon sequestration.

## Acknowledgements

GRC fieldwork and analysis was supported by a Peter Scott studentship at University of Cambridge. We are grateful to the Royal Society’s Southeast Asian Rainforest Research Programme (SEARRP) for logistical support and site access, Yayasan Sabah, Danum Valley Management Committee and the Sabah Biodiversity Council for permission to conduct research in Sabah, and to Toby Gardner, Daniel Field and Ben Phalan for useful discussions. For the purposes of open access, the author has applied a Creative Commons Attribution (CC BY) licence to any Author Accepted Manuscript version arising from this submission.

## Data availability

Code needed to replicate these analyses are provided at https://github.com/gcerullo/LianaBirdAnalysis. Data will be deposited on Zenodo upon manuscript review.

## References

1. Strassburg, B. B. N. et al. Global priority areas for ecosystem restoration. Nature 586, 724–729 (2020).

2. Edwards, D. P. et al. Upscaling tropical restoration to deliver environmental benefits and socially equitable outcomes. Curr Biol 31, R1326–R1341 (2021).

3. Pillay, R. et al. The Kunming-Montreal Global Biodiversity Framework needs headline indicators that can actually monitor forest integrity. ENVIRONMENTAL RESEARCH: ECOLOGY 3, (2024).

4. Cook-Patton, S. C. et al. Mapping carbon accumulation potential from global natural forest regrowth. Nature 585, 545–550 (2020).

5. Williams, B. A. et al. Global potential for natural regeneration in deforested tropical regions. Nature 636, 131–137 (2024).

6. Balmford, A. et al. Time to fix the biodiversity leak. Science 387, 720–722 (2025).

7. Cerullo, G. R. & Edwards, D. P. Actively restoring resilience in selectively logged tropical forests. Journal of Applied Ecology 56, 107–118 (2019).

8. Robinson, N. et al. Protect young secondary forests for optimum carbon removal. Preprint at 10.21203/rs.3.rs-4659226/v1 (2024).

9. Mo, L. et al. Integrated global assessment of the natural forest carbon potential. Nature 624, 92–101 (2023).

10. Bourgoin, C. et al. Human degradation of tropical moist forests is greater than previously estimated. Nature 631, 570–576 (2024).

11. Ewers, R. M. et al. Thresholds for adding degraded tropical forest to the conservation estate. Nature 631, 808–813 (2024).

12. Philipson, C. D. et al. Active restoration accelerates the carbon recovery of human-modified tropical forests. Science 369, 838–841 (2020).

13. Finlayson, C., Roopsind, A., Griscom, B. W., Edwards, D. P. & Freckleton, R. P. Removing climbers more than doubles tree growth and biomass in degraded tropical forests. Ecology and Evolution 12, e8758 (2022).

14. Putz, F. E. et al. Sustained timber yield claims, considerations, and tradeoffs for selectively logged forests. PNAS Nexus 1, pgac102 (2022).

15. Veryard, R. et al. Positive effects of tree diversity on tropical forest restoration in a field-scale experiment. Science Advances 9, eadf0938 (2023).

16. Putz, F. E. et al. Liana cutting in selectively logged forests increases both carbon sequestration and timber yields. Forest Ecology and Management 539, 121038 (2023).

17. Ruslandi Cropper, W. P. & Putz, F. E. Effects of silvicultural intensification on timber yields, carbon dynamics, and tree species composition in a dipterocarp forest in Kalimantan, Indonesia: An individual-tree-based model simulation. Forest Ecology and Management 390, 104–118 (2017).

18. Finlayson, C., Roopsind, A., Griscom, B. W., Edwards, D. P. & Freckleton, R. P. Removing climbers more than doubles tree growth and biomass in degraded tropical forests. Ecology and Evolution 12, e8758 (2022).

19. Cerullo, G. R., Edwards, F. A., Mills, S. C. & Edwards, D. P. Tropical forest subjected to intensive post-logging silviculture maintains functionally diverse dung beetle communities. Forest Ecology and Management 444, 318–326 (2019).

20. Soto-Navarro, C. et al. Mapping co-benefits for carbon storage and biodiversity to inform conservation policy and action. Phil. Trans. R. Soc. B 375, 20190128 (2020).

21. Griscom, B. W. et al. National mitigation potential from natural climate solutions in the tropics. Phil. Trans. R. Soc. B 375, 20190126 (2020).

22. Gilroy, J. J. et al. Cheap carbon and biodiversity co-benefits from forest regeneration in a hotspot of endemism. Nature Clim Change 4, 503–507 (2014).

23. Putz, F. E. et al. Intact Forest in Selective Logging Landscapes in the Tropics. Front. For. Glob. Change 2, (2019).

24. Miranda, E. B. P., Peres, C. A., Marini, M. Â. & Downs, C. T. Harpy Eagle (Harpia harpyja) nest tree selection: Selective logging in Amazon forest threatens Earth’s largest eagle. Biological Conservation 250, 108754 (2020).

25. Burivalova, Z. et al. Avian responses to selective logging shaped by species traits and logging practices. Proc Biol Sci 282, 20150164 (2015).

26. Lavery, T. H., Posala, C. K., Tasker, E. M. & Fisher, D. O. Ecological generalism and resilience of tropical island mammals to logging: A 23 year test. Global Change Biology 26, 3285–3293 (2020).

27. Ansell, F. A., Edwards, D. P. & Hamer, K. C. Rehabilitation of Logged Rain Forests: Avifaunal Composition, Habitat Structure, and Implications for Biodiversity-Friendly REDD+: Rain forest Rehabilitation and Avifaunal Composition. Biotropica 43, 504–511 (2011).

28. Cosset, C. C. P., Gilroy, J. J. & Edwards, D. P. Impacts of tropical forest disturbance on species vital rates. Conservation Biology 33, 66–75 (2019).

29. Hayward, R. M. et al. Three decades of post-logging tree community recovery in naturally regenerating and actively restored dipterocarp forest in Borneo. Forest Ecology and Management 488, 119036 (2021).

30. Reynolds, G., Payne, J., Sinun, W., Mosigil, G. & Walsh, R. P. D. Changes in forest land use and management in Sabah, Malaysian Borneo, 1990–2010, with a focus on the Danum Valley region. Philosophical Transactions of the Royal Society B: Biological Sciences 366, 3168–3176 (2011).

31. Fisher, B. et al. Cost-effective conservation: calculating biodiversity and logging trade-offs in Southeast Asia. Conservation Letters 4, 443–450 (2011).

32. Edwards, D. et al. Degraded lands worth protecting: the biological importance of Southeast Asia’s repeatedly logged forests. Proceedings of the Royal Society B: Biological Sciences 278, 82–90 (2011).

33. Mitchell, S. L. et al. Riparian reserves help protect forest bird communities in oil palm dominated landscapes. Journal of Applied Ecology 55, 2744–2755 (2018).

34. Socolar, J. B., Mills, S. C., Haugaasen, T., Gilroy, J. J. & Edwards, D. P. Biogeographic multi-species occupancy models for large-scale survey data. Ecology and Evolution 12, e9328 (2022).

35. Socolar, J. B. & Mills, S. C. Introducing Flocker: An R Package for Flexible Occupancy Modeling via Brms and Stan. http://biorxiv.org/lookup/doi/10.1101/2023.10.26.564080 (2023) doi:10.1101/2023.10.26.564080.

36. Mills, S. C. et al. High sensitivity of tropical forest birds to deforestation at lower altitudes. Ecology 104, e3867 (2023).

37. Banks-Leite, C. et al. Assessing the utility of statistical adjustments for imperfect detection in tropical conservation science. Journal of Applied Ecology 51, 849–859 (2014).

38. Asner, G. P. et al. Mapped aboveground carbon stocks to advance forest conservation and recovery in Malaysian Borneo. Biological Conservation 217, 289–310 (2018).

39. Makowski, D., Ben-Shachar, M. S., Chen, S. H. A. & Lüdecke, D. Indices of Effect Existence and Significance in the Bayesian Framework. Front. Psychol. 10, (2019).

40. Cerullo, G. et al. Sparing old-growth maximises conservation outcomes within selectively logged Amazonian rainforest. Biological Conservation 282, 110065 (2023).

41. Burivalova, Z., Sekercioglu, Ç. H. & Koh, L. P. Thresholds of Logging Intensity to Maintain Tropical Forest Biodiversity. Current Biology 24, 1893–1898 (2014).

42. Matos, F. A. R. et al. Secondary forest fragments offer important carbon and biodiversity cobenefits. Global Change Biology 26, 509–522 (2020).

43. Seddon, N. et al. Getting the message right on nature-based solutions to climate change. Global Change Biology 27, 1518–1546 (2021).

44. Putz, F. E. et al. Sustained timber yield claims, considerations, and tradeoffs for selectively logged forests. PNAS Nexus 1, pgac102 (2022).

45. Campbell, M. J., Edwards, W., Odell, E., Mohandass, D. & Laurance, W. F. Can Lianas Assist in Rainforest Restoration? Tropical Conservation Science 8, 257–273 (2015).

46. Velho, N., Ratnam, J., Srinivasan, U. & Sankaran, M. Shifts in community structure of tropical trees and avian frugivores in forests recovering from past logging. Biological Conservation 153, 32–40 (2012).

47. Fricke, E. C., Cook-Patton, S. C., Harvey, C. F. & Terrer, C. Seed dispersal disruption limits tropical forest regrowth. Proceedings of the National Academy of Sciences 122, e2500951122 (2025).

48. Cosset, C. C. P. & Edwards, D. P. The effects of restoring logged tropical forests on avian phylogenetic and functional diversity. Ecological Applications 27, 1932–1945 (2017).

49. Johnson, O. et al. Amazonian birds in more dynamic habitats have less population genetic structure and higher gene flow. Molecular Ecology 32, 2186–2205 (2023).

50. França, F. et al. Do space-for-time assessments underestimate the impacts of logging on tropical biodiversity? An Amazonian case study using dung beetles. Journal of Applied Ecology 53, 1098–1105 (2016).

51. França, F. M. et al. Selective logging intensity and time since logging drive tropical bird and dung beetle diversity: a case study from Amazonia. Environmental Conservation 51, 112–121 (2024).

52. Marshall, A. R. et al. Conceptualising the Global Forest Response to Liana Proliferation. Front. For. Glob. Change 3, (2020).

53. Ngute, A. S. K. et al. Global dominance of lianas over trees is driven by forest disturbance, climate and topography. Global Change Biology 30, e17140 (2024).

54. Harrison, R. D. et al. Restoration concessions: a second lease on life for beleaguered tropical forests? Frontiers in Ecology and the Environment 18, 567–575 (2020).

55. Sagar, H. S. S. C. et al. Avifauna recovers faster in areas less accessible to trapping in regenerating tropical forests. Biological Conservation 279, 109901 (2023).

56. Morton, O., Scheffers, B. R., Haugaasen, T. & Edwards, D. P. Impacts of wildlife trade on terrestrial biodiversity. Nature Ecology & Evolution 5, 540–548 (2021).

57. Nishijima, S. et al. Evaluating the impacts of wood production and trade on bird extinction risks. Ecological Indicators 71, 368–376 (2016).

58. Burivalova, Z. et al. Does biodiversity benefit when the logging stops? An analysis of conservation risks and opportunities in active versus inactive logging concessions in Borneo. Biological Conservation 241, 108369 (2020).

59. Milodowski, D. T. et al. The impact of logging on vertical canopy structure across a gradient of tropical forest degradation intensity in Borneo. Journal of Applied Ecology 58, 1764–1775 (2021).

60. Malhi, Y. et al. Logged tropical forests have amplified and diverse ecosystem energetics. Nature 612, 707–713 (2022).

