## Supplementary figures for "Persistent declines in forest-dependent birds following active restoration of logged tropical forest in Borneo"

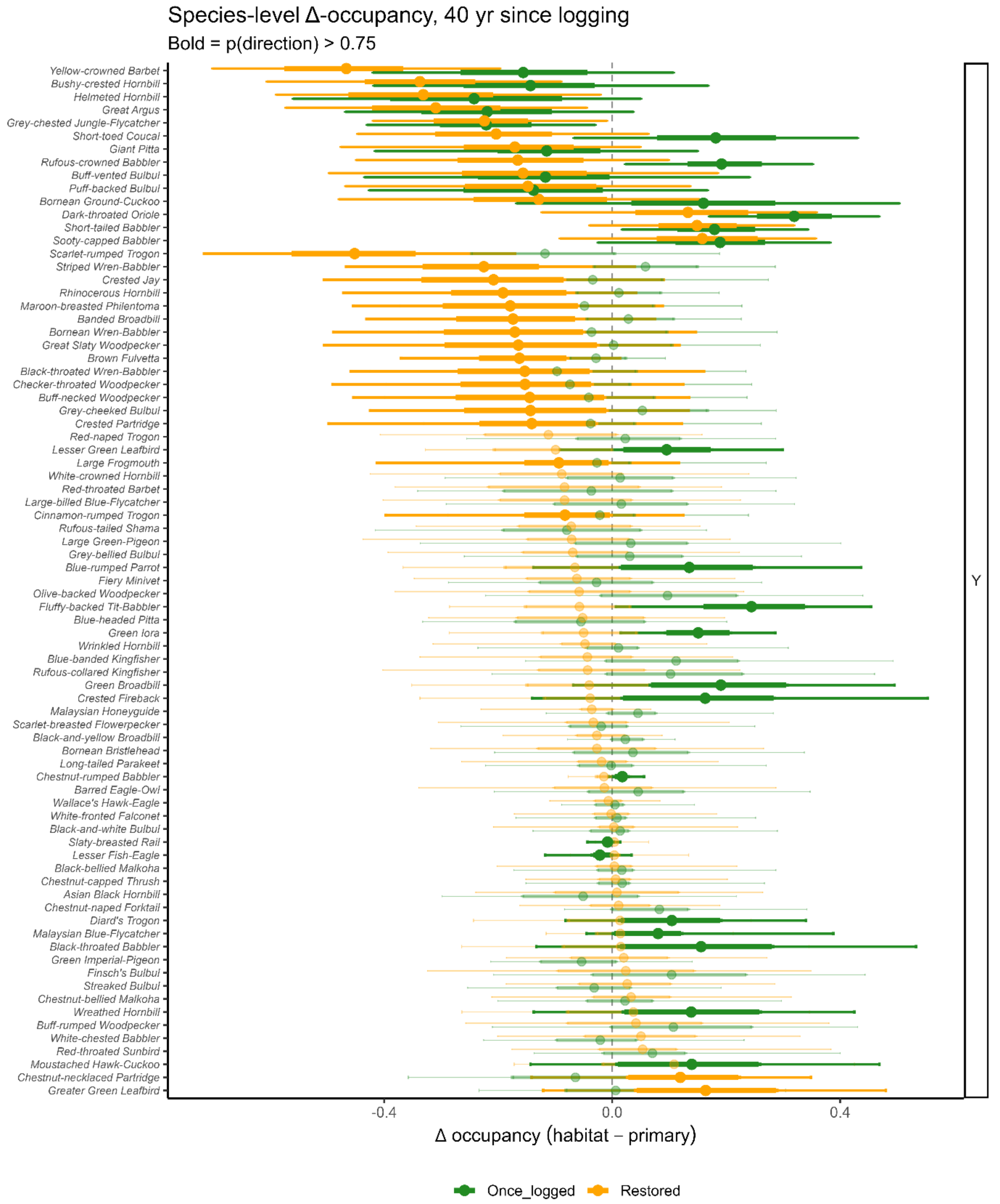


**Figure S1.** Threatened and near-threatened species-level changes in occupancy in once-logged and restored forests relative to primary forest, across a lowland dipterocarp rainforest landscape in Sabah, Borneo. Each point represents the mean difference in occupancy (Δ occupancy = habitat – primary) for a species in once-logged (green) and restored (orange) forests, estimated at 40 years post-logging. Thick lines indicate 50% credible intervals and thin lines indicate 90% credible intervals, calculated from 500 posterior draws per species. Species are ordered by mean Δ occupancy in restored forests. The vertical dashed line at zero indicates no difference in occupancy between disturbed and primary forest. Positive values reflect higher occupancy in once-logged or restored forest than in primary forest, and negative values reflect lower occupancy. Points shown in bold have a probability of direction (pd) > 0.75, suggesting that the estimated difference from primary forest is relatively consistent across posterior draws and therefore likely reflects a genuine directional effect rather than random variation.

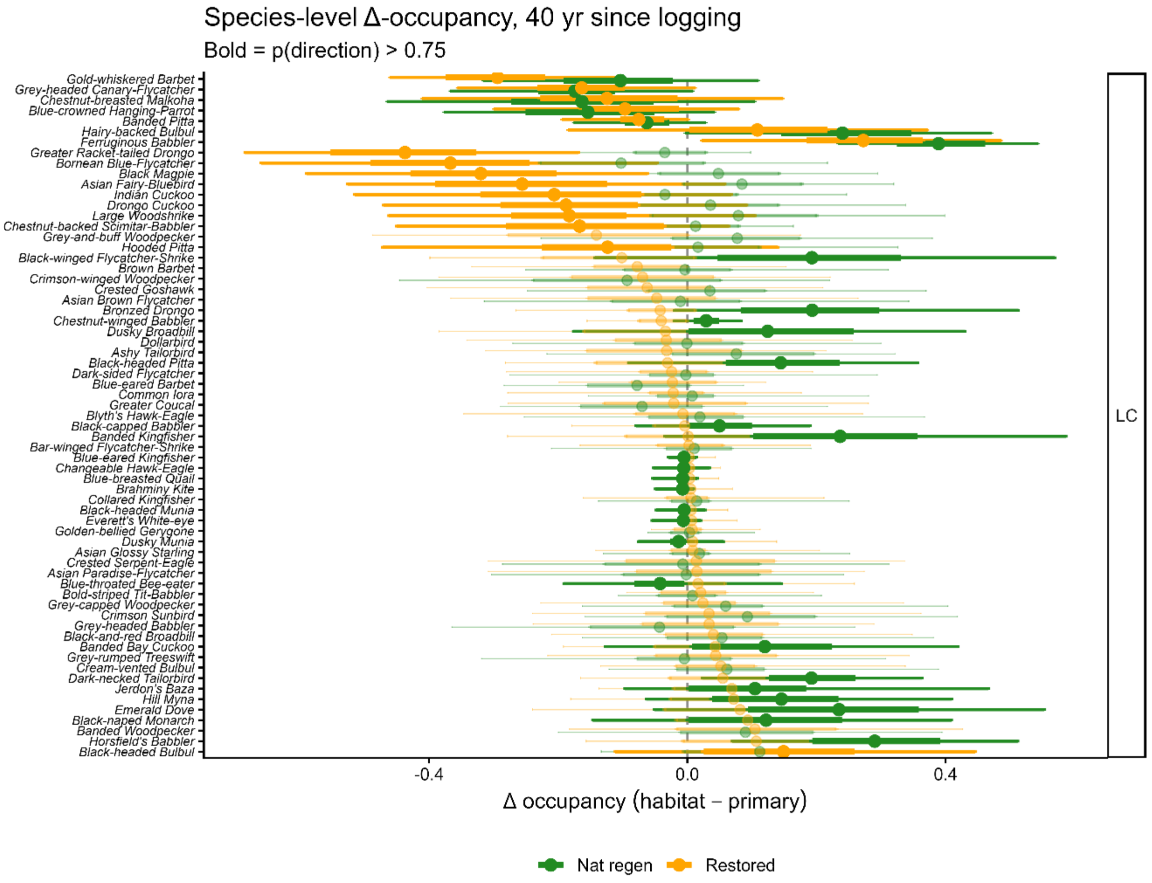

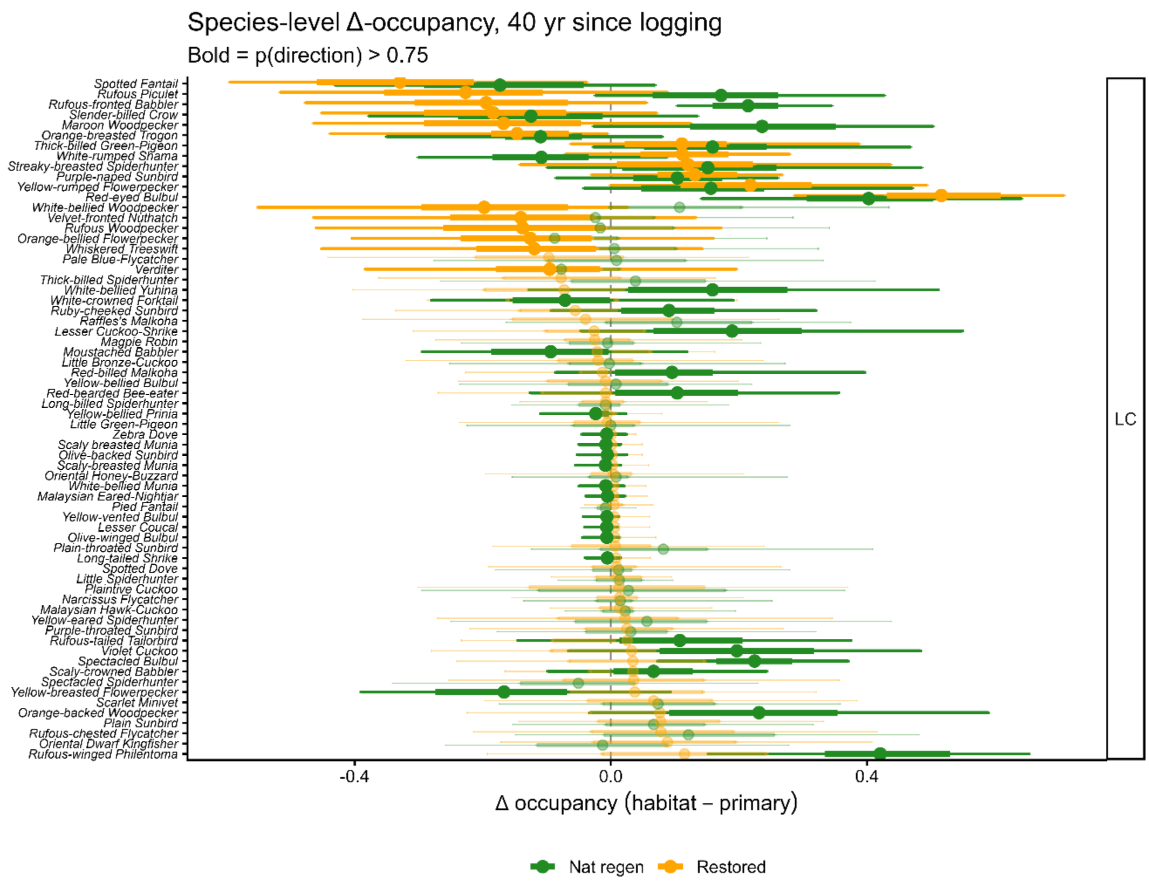


**Figure S2.** Same as Figure S1 but for species listed by the IUCN as of global "Least concern".

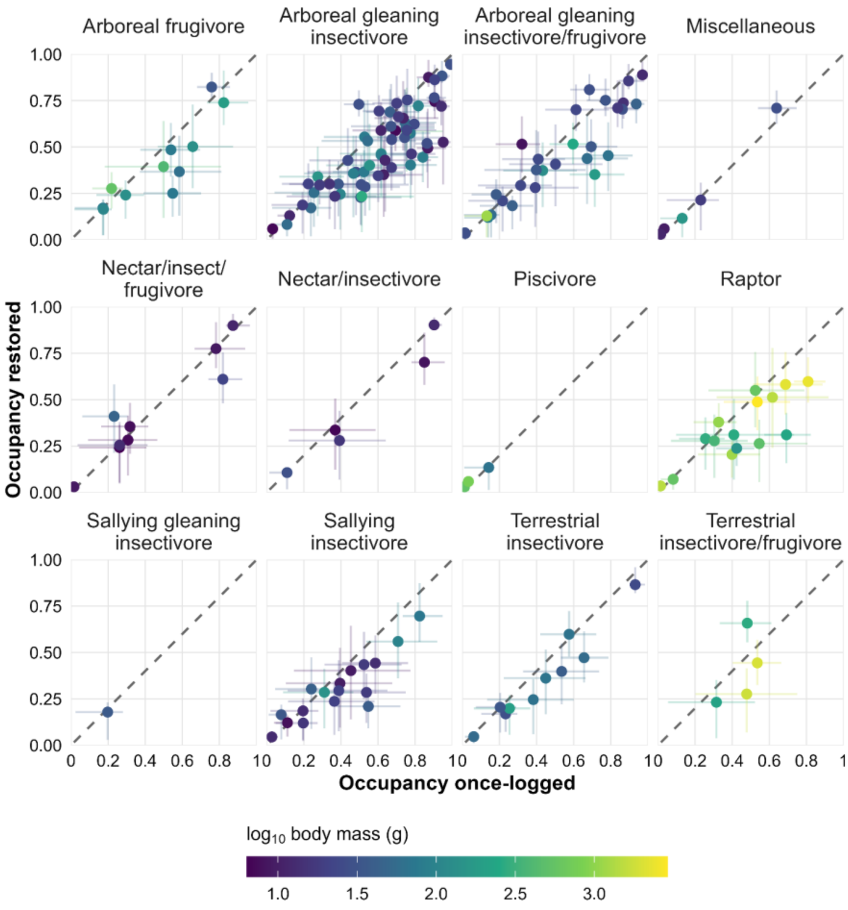


**Figure S3.** Species responses to restoration interventions across feeding-foraging guilds, with species body mass (log10) showing the relative masses of different species.

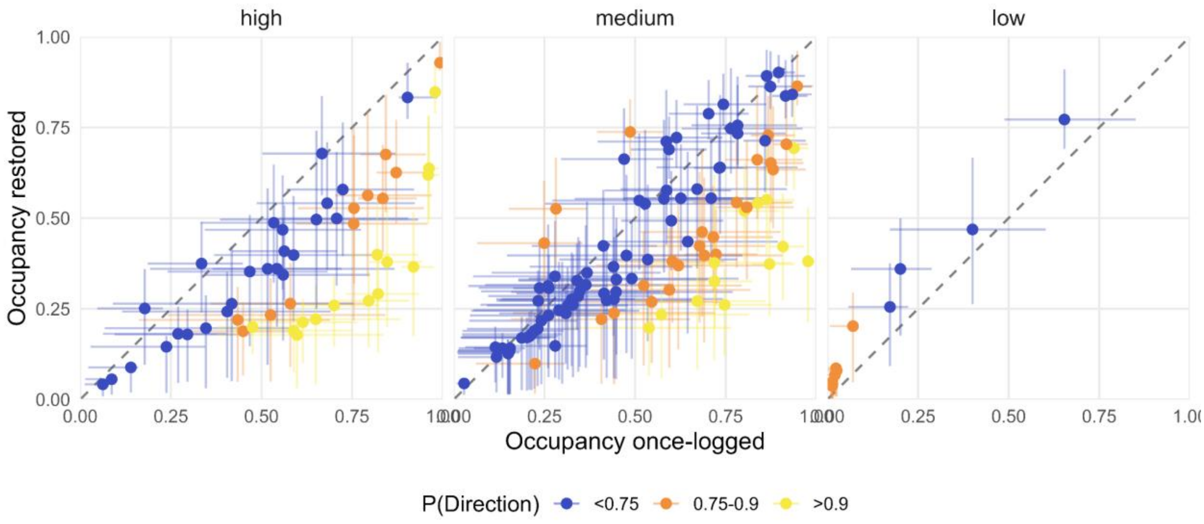


**Figure S4**. Same as Figure 2 & 3 but for forest dependency

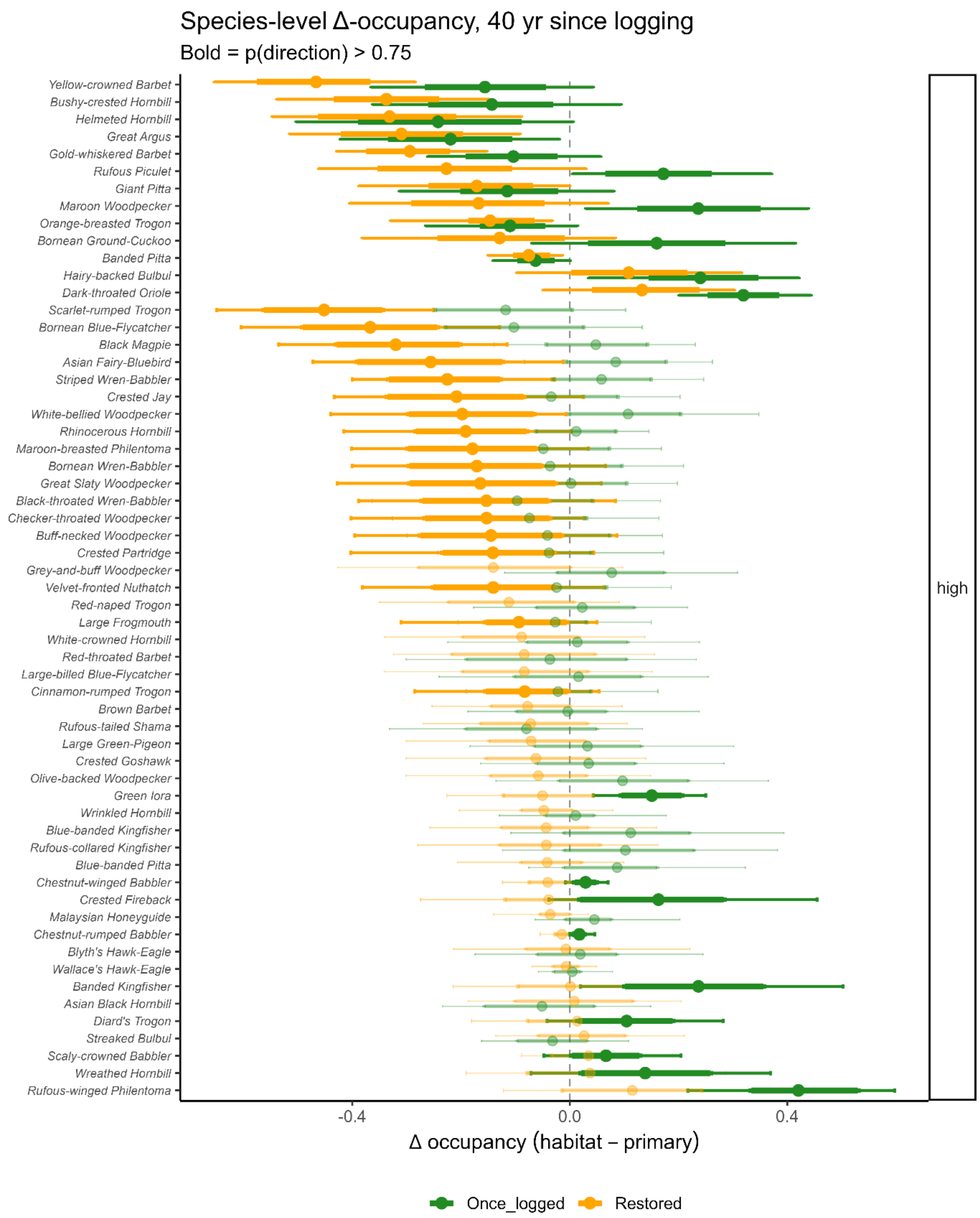


**Figure S5.** Same as Figure S1 but for high-dependency birds 40 yrs after logging.

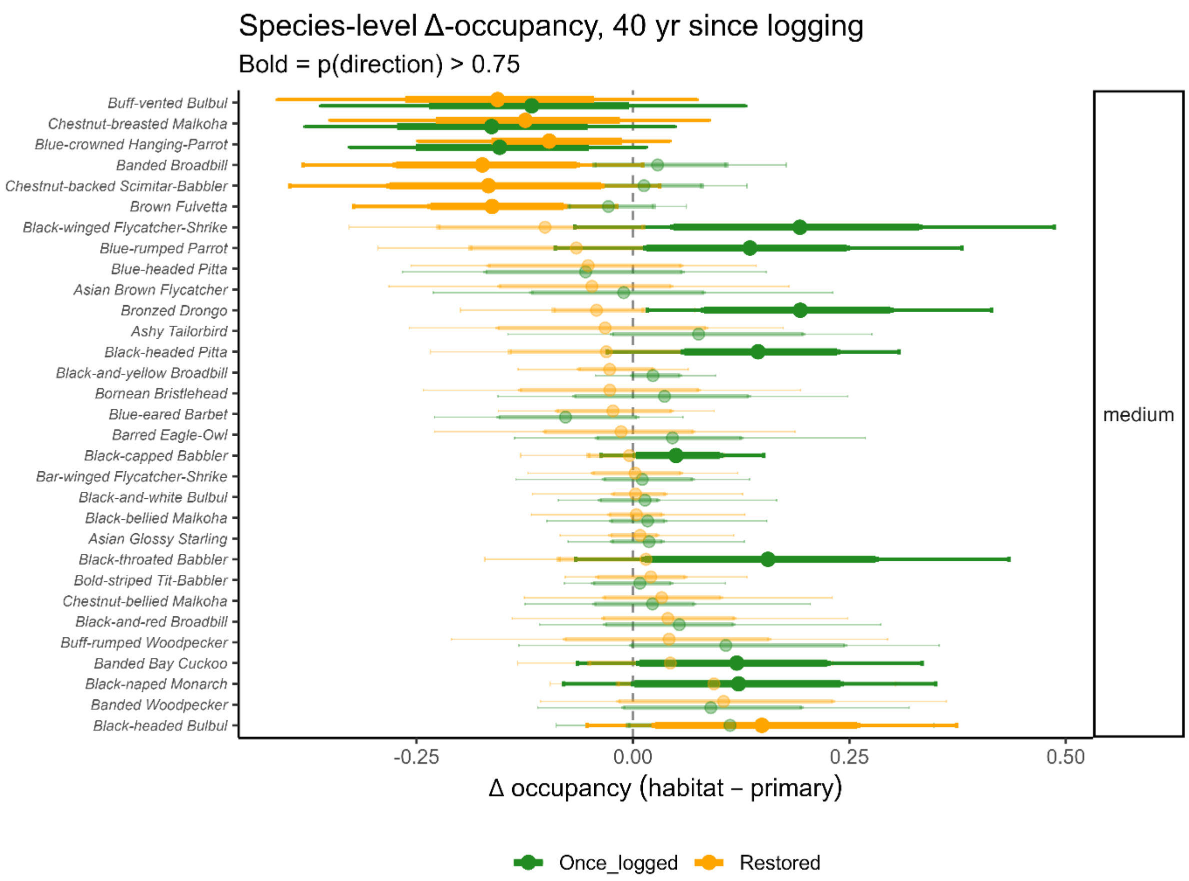
Species-level Δ-occupancy, 40 yr since logging
Bold = p(direction) > 0.75
Buff-vented Bulbul
Chestnut-breasted Malkoha
Blue-crowned Hanging-Parrot
Banded Broadbill
Chestnut-backed Scimitar-Babbler
Brown Fulvetta
Black-winged Flycatcher-Shrike
Blue-rumped Parrot
Blue-headed Pitta
Asian Brown Flycatcher
Bronzed Drongo
Ashy Tailorbird
Black-headed Pitta
Black-and-yellow Broadbill
Bornean Bristlehead
Blue-eared Barbet
Barred Eagle-Owl
Black-capped Babbler
Bar-winged Flycatcher-Shrike
Black-and-white Bulbul
Black-bellied Malkoha
Asian Glossy Starling
Black-throated Babbler
Bold-striped Tit-Babbler
Chestnut-bellied Malkoha
Black-and-red Broadbill
Buff-rumped Woodpecker
Banded Bay Cuckoo
Black-naped Monarch
Banded Woodpecker
Black-headed Bulbul
medium
-0.25
0.00
0.25
0.50
Δ occupancy (habitat – primary)
Once_logged
Restored

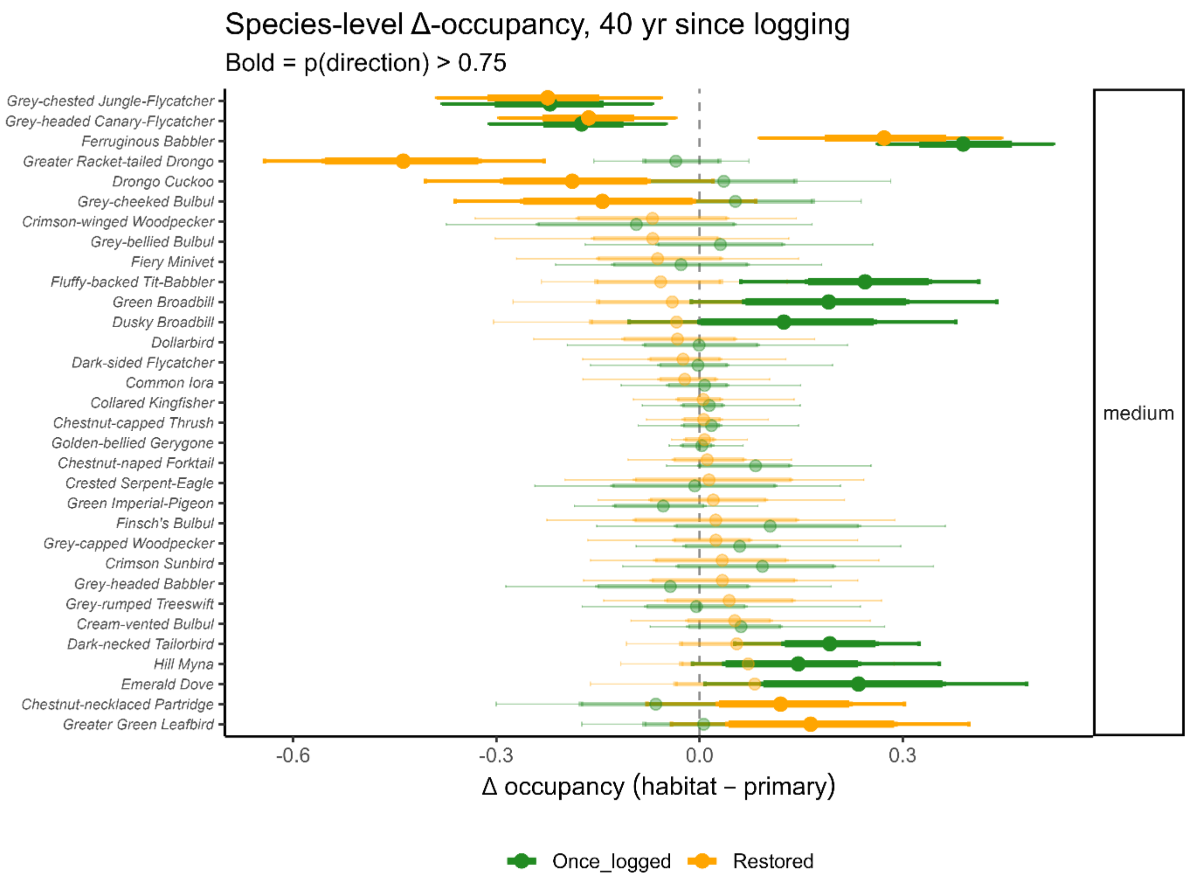
Species-level Δ-occupancy, 40 yr since logging
Bold = p(direction) > 0.75
Grey-chested Jungle-Flycatcher
Grey-headed Canary-Flycatcher
Ferruginous Babbler
Greater Racket-tailed Drongo
Drongo Cuckoo
Grey-cheeked Bulbul
Crimson-winged Woodpecker
Grey-bellied Bulbul
Fiery Minivet
Fluffy-backed Tit-Babbler
Green Broadbill
Dusky Broadbill
Dollarbird
Dark-sided Flycatcher
Common Iora
Collared Kingfisher
Chestnut-capped Thrush
Golden-bellied Gerygone
Chestnut-naped Forktail
Crested Serpent-Eagle
Green Imperial-Pigeon
Finsch's Bulbul
Grey-capped Woodpecker
Crimson Sunbird
Grey-headed Babbler
Grey-rumped Treeswift
Cream-vented Bulbul
Dark-necked Tailorbird
Hill Myna
Emerald Dove
Chestnut-necklaced Partridge
Greater Green Leafbird
medium
-0.6
-0.3
0.0
0.3
Δ occupancy (habitat – primary)
Once_logged
Restored

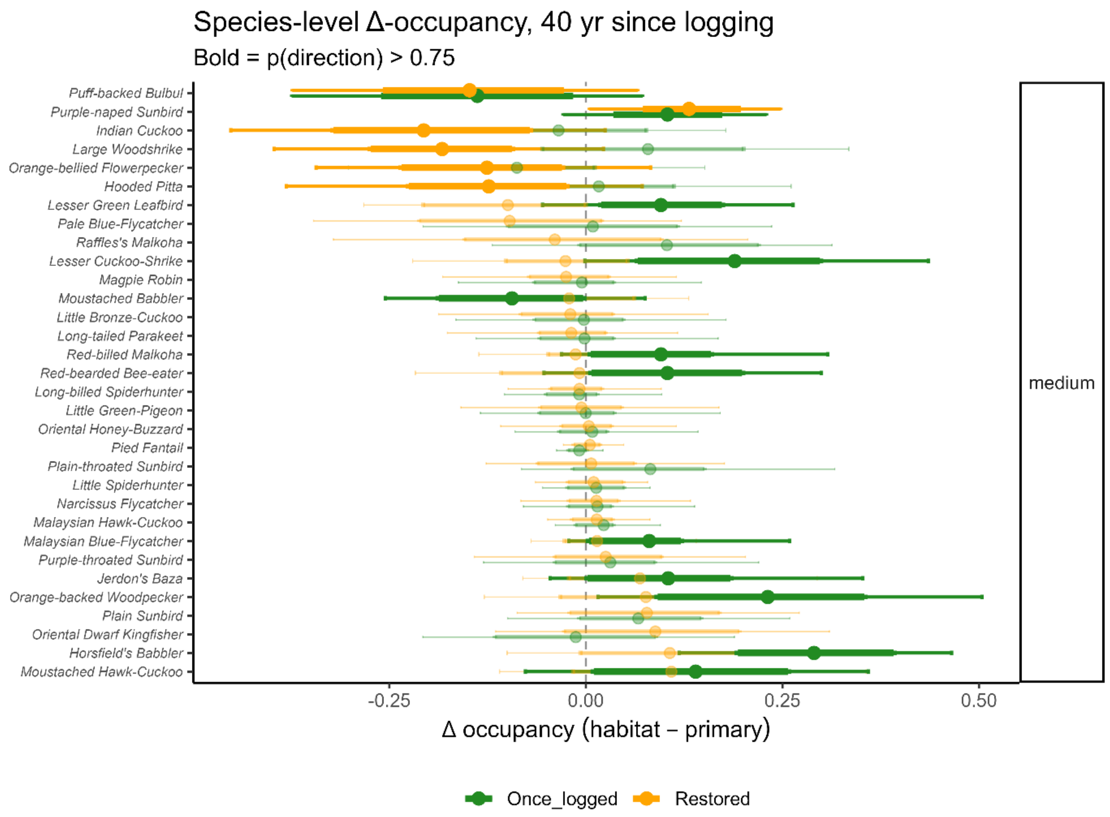


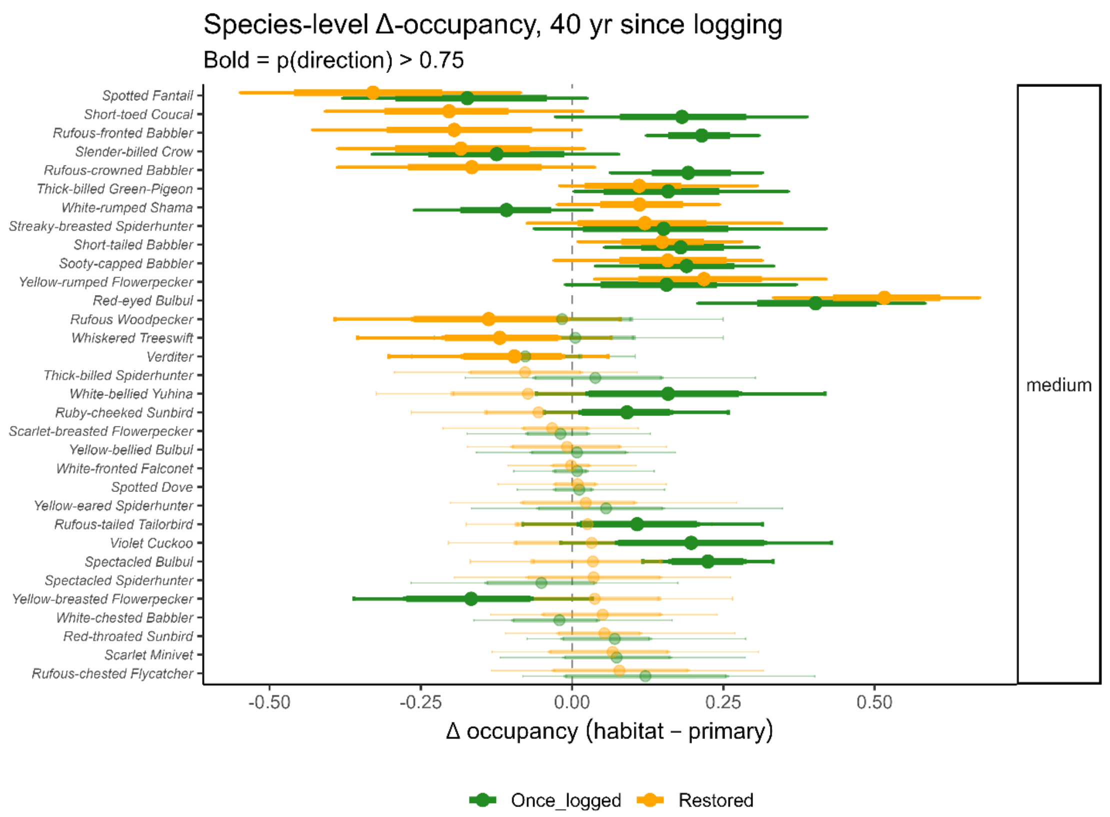


**Figure S6.** Same as Figure S1 but for medium-dependency birds 40-yrs after logging.

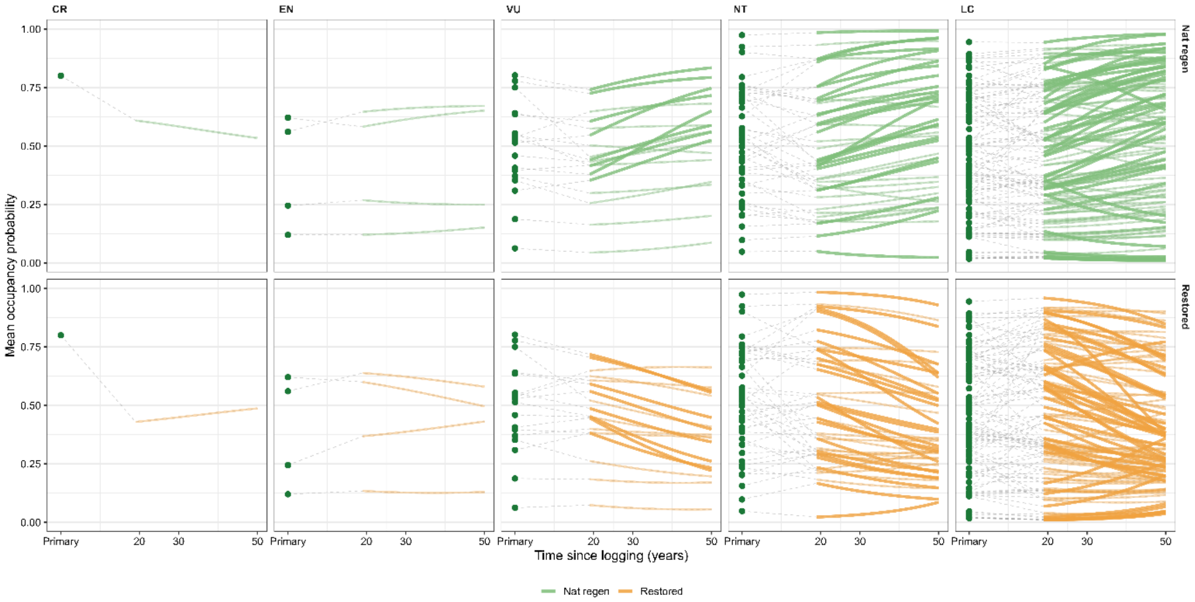


**Figure S7**. Same as Figure 4 but by IUCN threat category.

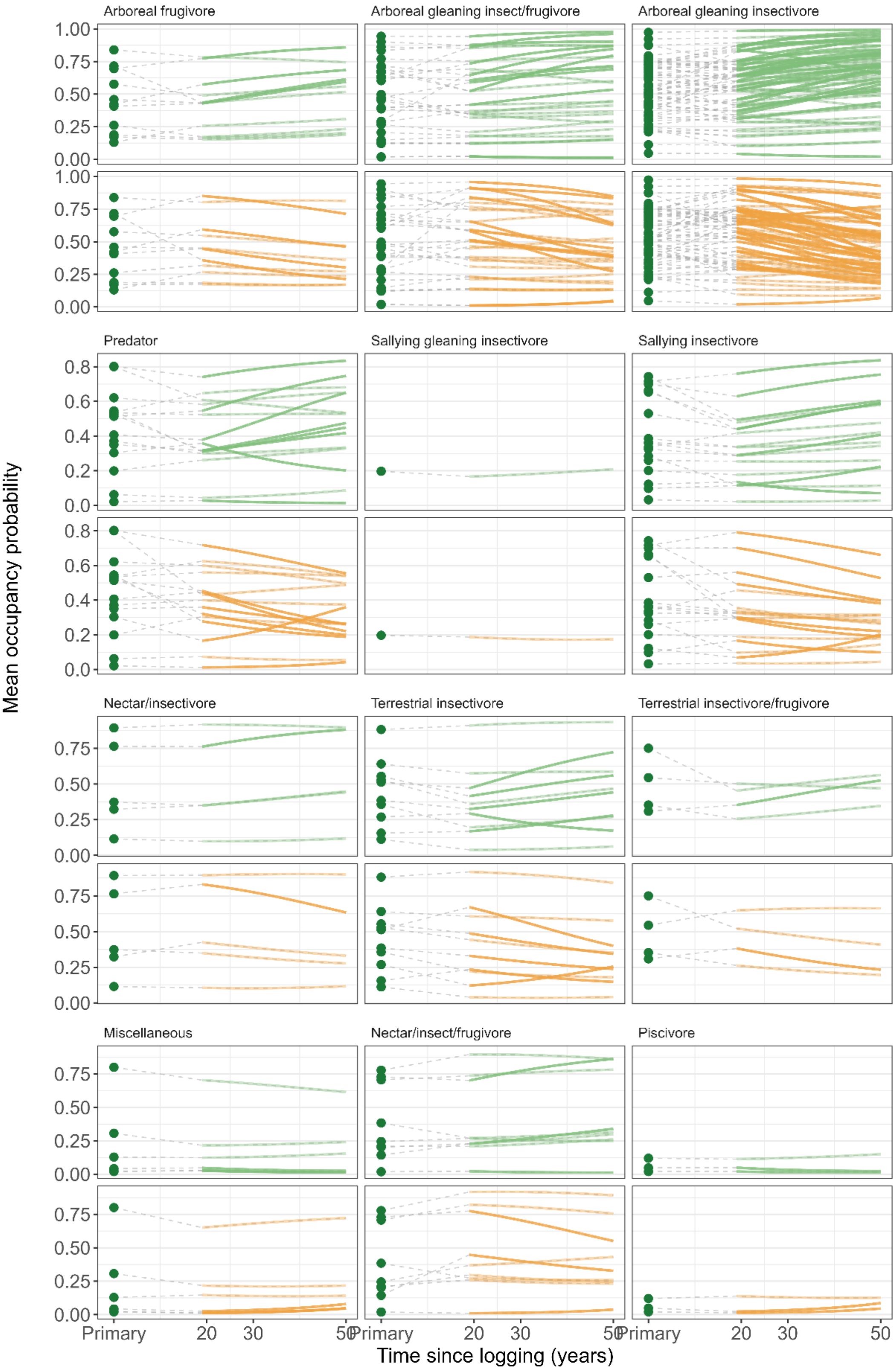

Arboreal frugivore
Arboreal gleaning insect/frugivore
Arboreal gleaning insectivore
Predator
Sallying gleaning insectivore
Sallying insectivore
Nectar/insectivore
Terrestrial insectivore
Terrestrial insectivore/frugivore
Miscellaneous
Nectar/insect/frugivore
Piscivore
Mean occupancy probability
Time since logging (years)
Primary
20
30
50
0.00
0.25
0.50
0.75
1.00
0.0
0.2
0.4
0.6
0.8

**Figure S8**. Same as Figure 4 but for species feeding-foraging guild.
